# Zinc-binding to the cytoplasmic PAS domain regulates the essential WalK histidine kinase of *Staphylococcus aureus*

**DOI:** 10.1101/405365

**Authors:** Ian R. Monk, Nausad Shaikh, Stephanie L. Begg, Mike Gajdiss, Jean Y. H. Lee, Sacha J. Pidot, Torsten Seemann, Michael Kuiper, Brit Winnen, Rikki Hvorup, Brett M. Collins, Gabriele Bierbaum, Benjamin P. Howden, Christopher A. McDevitt, Glenn F. King, Timothy P. Stinear

**Author notes:** Contributed equally (Ian R. Monk, Nausad Shaikh).

## Abstract

WalKR (YycFG) is the only essential two-component regulator in the human pathogen *Staphylococcus aureus.* WalKR regulates peptidoglycan synthesis, but this function alone appears not to explain its essentiality. To understand WalKR function we investigated a suppressor mutant that arose when WalKR activity was impaired; a single histidine to tryptophan substitution (H271Y) in the cytoplasmic Per-Arnt-Sim (PAS^CYT^) domain of the histidine kinase WalK. Introduction of the WalK^H271Y^ mutation into wild-type *S*. *aureus* activated the WalKR regulon. Structural analyses of the WalK PAS^CYT^ domain revealed a hitherto unknown metal binding site, in which a zinc ion (Zn^2+^) was tetrahedrally-coordinated by four amino acid residues including H271. The Wallk^H271Y^ mutation abrogated metal binding, increasing WalK kinase activity and WalR phosphorylation. Thus, Zn^2+^-binding negatively regulates WalKR activity. Identification of a metal ligand sensed by the WalKR system substantially expands our understanding of this critical *S*. *aureus* regulon.

## Introduction

*Staphylococcus aureus* is a major human pathogen that causes a wide range of hospital- and community-acquired opportunistic infections^1^ Antibiotic-resistant strains (in particular methicillin-resistant *S*. *aureus* [MRSA]) are increasing in prevalence in both hospital and community settings. Resistance to last line agents such as vancomycin, linezolid and daptomycin is well described ^23^, casting doubt on their efficacy for the treatment of serious MRSA infections ^45^. In the context of limited treatment options, the social and financial burden posed by *S*. *aureus* related disease is now globally significant^1^

WalKR is a highly conserved two-component regulatory system (TCS) with features that are unique among low G+C Gram-positive bacteria^6^. Like other TCS, it comprises a multi-domain transmembrane sensor histidine kinase (WalK – HK) (Fig. 1a) and a response regulator (WalR – RR). Notably, in *S*. *aureus* and closely related bacteria, the locus is essential, a characteristic that has made it a potential therapeutic target^7^. The WalR/WalK system (also called YycFG and VicRK) was first identified from temperature-sensitive mutants in *Bacillus subtilis* ^8^ and then *S*. *aureus* ^9^, with the system essential in both genera. Essentiality of WalKR has been inferred through both construction of strains containing an inducible WalKR ^10,11^, which were unable to grow in the absence of inducer, and the inability to delete the genes in *Listeria monocytogenes* or *Enterococcus faecalis* ^12,13^. In an impressive recent study, Villanueva *et al*^14^. deleted all 15 non-essential TCSs from two differents. *aureus* backgrounds and confirmed that WalKR is the only TCS strictly required for growth. Depletion of WalKR in *B. subtilis* produces a long-chain phenotype with the formation of multiple ‘ghost’ cells without cytoplasmic contents, which correlates with a reduction in colony forming units (CFU)^15^. In *S*. *aureus,* depletion of WalKR causes the same loss in viability, but without cellular lysis, although aberrant septum formation was reported ^16^. These observations suggest different mechanisms for the essentiality of WalKR in rod versus coccus shaped bacteria. However, this is not a uniform requirement as although the TCS is also found in streptococci, the WalK ortholog in this species is not essential and downstream regulators are absent, suggesting altered regulation. Further, WalK in streptococci contain only one transmembrane domain, which differentiates it from other cocci^6^.

**Figure 1.**
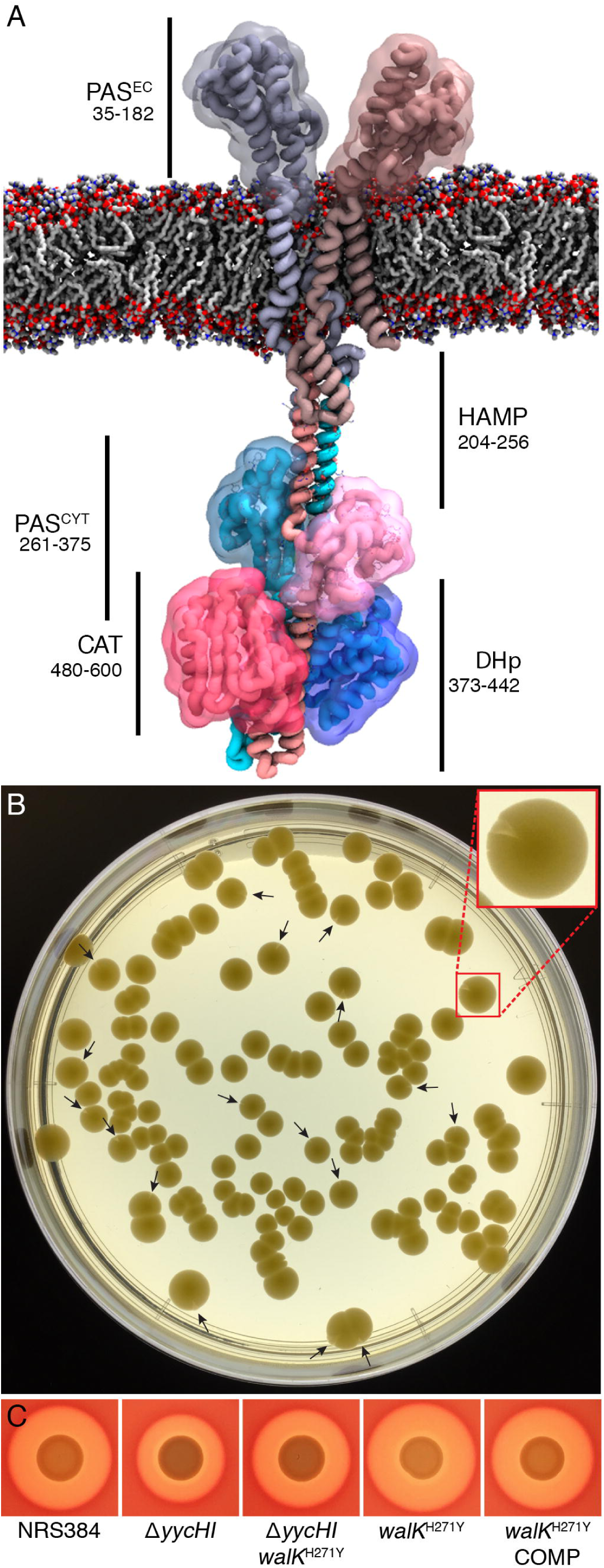
Colony sectoring in aΔyycHI leads to mutation in WalK. **(A)** Molecular model of the essential two component histidine kinase WalK (dimer 608 residues) from *S. aureus* in a phospholipid bilayer. The amino acid boundaries of the various WalK domains (PAS^EC^: Extracellular PAS; PAS^CYT^: Cytoplasmic PAS; DHp: Dimerization and Histidine Phosphorylation; CAT: Catalytic/ATP binding) are highlighted. **(B)** Plating of NRS384ΔyycA7/ onto BHI agar after 48 h at 37°C promotes colony sectoring (arrow heads). Red box inset shows enlarged view of one sectored colony. **(C)** The WalK^H271Y^ mutation identified from whole genome sequencing of a single sectored colony was introduced by allelic exchange into the NRS384 background, with the mutation increasing hemolysis on sheep blood agar. Also shown are the wild-type (NRS384), Δ*yycH, ΔyycHI walK* ^H271Y^and the m/o/K^H271Y-COMP^, for reference.

S. *aureus antibiotic* resistance and clinical persistence are frequently linked to mutations in regulatory genes, in particular TCS. Among the *S. aureus* TCSs, mutations in loci such as *vraRS, graRS* and *walKR* a reassociated with the vancomycin intermediate resistant *S. aureus* (VISA) phenotype ^17^-^19^. Notably, mutations in *WalKR* are a critical contributor toward this phenotype with numerous clinical VISA strains possessing *walKR* mutations ^20^. Induction of characteristic VISA phenotypes (thickened cell wall, reduced autolytic activity and reduced virulence) can arise from mutations as simple as a single nucleotide change in either *walK* or *walR* ^21^-^23^.

Despite the central role of the TCS in bacterial viability, the physiological and/or mechanical signal sensed by WalK is unknown. In *B. subtilis,* where it has been more extensively studied, WalK localises to the division septum and interacts with proteins of the divisome ^24^. *B. subtilis* WalR positively regulates several autolysins involved in peptidoglycan metabolism and represses inhibitors of these enzymes ^25^. Thus, WalKR has been inferred to link cell division with cell wall homeostasis ^624^. Nevertheless, its role in *S*. *aureus* appears to be distinct. Although *S*. *aureus* WalR does control autolysin expression ^16,26^, this function does not explain the essentiality of the system as expression of genes encoding lytic transglycosylases or amidases in a WalKR-depleted strain do not restore cell viability ^27^. Further, the membrane associated regulators YycHI act as an activator of WalK function in *S*. *aureus* ^*28*^, which contrasts with *B. subtilis* where they serve to repress the system ^29^.

Sequence variation provides one basis for the apparent differences in WalKR function between *S*. *aureus* and *B. subtilis.* The WalK alleles share only 45% amino acid identity, with the majority of the variation concentrated in the extracellular region and a cytoplasmic Per-Arnt Sim (PAS) domain (Fig. 1a). Although these regions have low sequence conservation, PAS domains are known to adopt a conserved tertiary fold, where they facilitate sensory perception via ligand interaction enabling signal transduction. In HKs that contain PAS domains, known ligands include heme, flavin mononucleotide and di/tricarboxylic acids^30^. In *B. subtilis,* the cytoplasmic PAS domain is essential for WalK localisation to the division septum ^24^. Recently, *S*. *aureus* WalK was also shown via a GFP-fusion to be to be preferentially located to the division septum. The role of the cytoplasmic PAS domain in septum targeting was not examined^31^.

Here, we screened *S*. *aureus yycHI* deletion mutants and identified a novel WalK suppressor mutation (H271Y) located in the cytoplasmic PAS domain. Structural and functional analyses of WalK revealed that this residue is a critical component of a cytoplasmic metal binding site that directly influenced HK activity. This metalloregulatory site was observed to bind zinc (Zn^2+^) *in vitro,* and this was abrogated by the H271Y mutation resulting in increased WalK autophosphorylation. These observations were corroborated *in vivo* through the construction of a metal-binding site mutant at the native locus with subsequent activation of WalKR-associated phenotypes and increased WalR phosphorylation *in vivo.* Molecular modelling of WalK in Zn^2+^-free and Zn^2+^-bound states suggests that metal occupancy influences conformational changes associated with the cytoplasmic domains, thus providing a plausible mechanism of activity modulation. To our knowledge, this is the first description of metal binding to the cytoplasmic PAS domain influencing the activity of a histidine kinase.

## RESULTS

### Mapping of a suppressor mutation to *walK* by genome sequencing

YycH and Yycl are membrane-associated accessory proteins that have been shown to directly interact with WalK and positively regulate WalKR function ^28^. We constructed an *S*. *aureus yycHI* deletion mutant (*ΔyycHi)* in USA300 strain NRS384 and observed colony sectoring after 2 days growth at 37°C on BHI agar (Fig. 1b), suggesting the development of suppressor (compensatory) mutations. We purified the two colony morphotypes and performed whole genome sequencing. Aligning the sequence reads to the closed NRS384 genome revealed only a single point mutation in addition to the engineered *yycHI* deletion. The mutation occurred in the cytoplasmic PAS domain of WalK, wherein histidine 271 was predicted to be replaced by tyrosine (WalK^H271Y^). Despite the different chemical profiles of the two side chains, *i.e.* positively charged versus slightly polar, the two residues have similar steric bulk.

### Generation of a *walk* ^H271Y^ site-directed mutant

We next sought to investigate the impact of the WalK^H271Y^ substitution by introducing the mutation into wild-type NRS384 by allelic exchange. The resulting unmarked mutant was analysed by genome sequencing to exclude the occurrence of secondary site mutations, as was previously observed to occur in backgrounds involving mutagenesis of WalK ^22,32^. Here, only the nucleotides targeted by the allelic exchange of *walK* differed between wild-type *walK* and the *walK*^*H271Y*^ mutant. The allelic exchange procedure was then repeated to convert the *walK* ^H*271Y*^ allele back to wild-type with the introduction of a silent *Pst\* restriction site to mark the revertant *(walK*^*H271Y-COMP*^). By comparison with the wild-type, the *wak* ^H*271Y*^ mutant formed smaller colonies on sheep blood agar with reduced pigmentation, but produced a slightly larger zone of hemolysis (Fig. 1c). Deletion of *yycHI* resulted in reduced hemolysis. However, hemolysis was elevated upon the introduction of the *walK* ^H*271Y*^ allele into the Δ*yycHI* background (Fig. 1c). Intriguingly, following growth in TSB at 37°C, the WalK^H271Y^strain did not exhibit a lag phase upon inoculation into fresh medium (Fig. 2a). Nonetheless, at 2 h post inoculation the growth rate was significantly reduced in comparison with the wild-type and complemented strain. The maximal doubling rate was 33 min for the WalK^H271Y^ mutant versus 23 min for the wild-type and complemented strains. Further, the WalK^H271Y^ mutant strain had a reduced final optical density at 600 nm (OD_600_) compared to the wild-type, although this equated to identical CFU counts (Fig. 2a).

**Figure 2.**
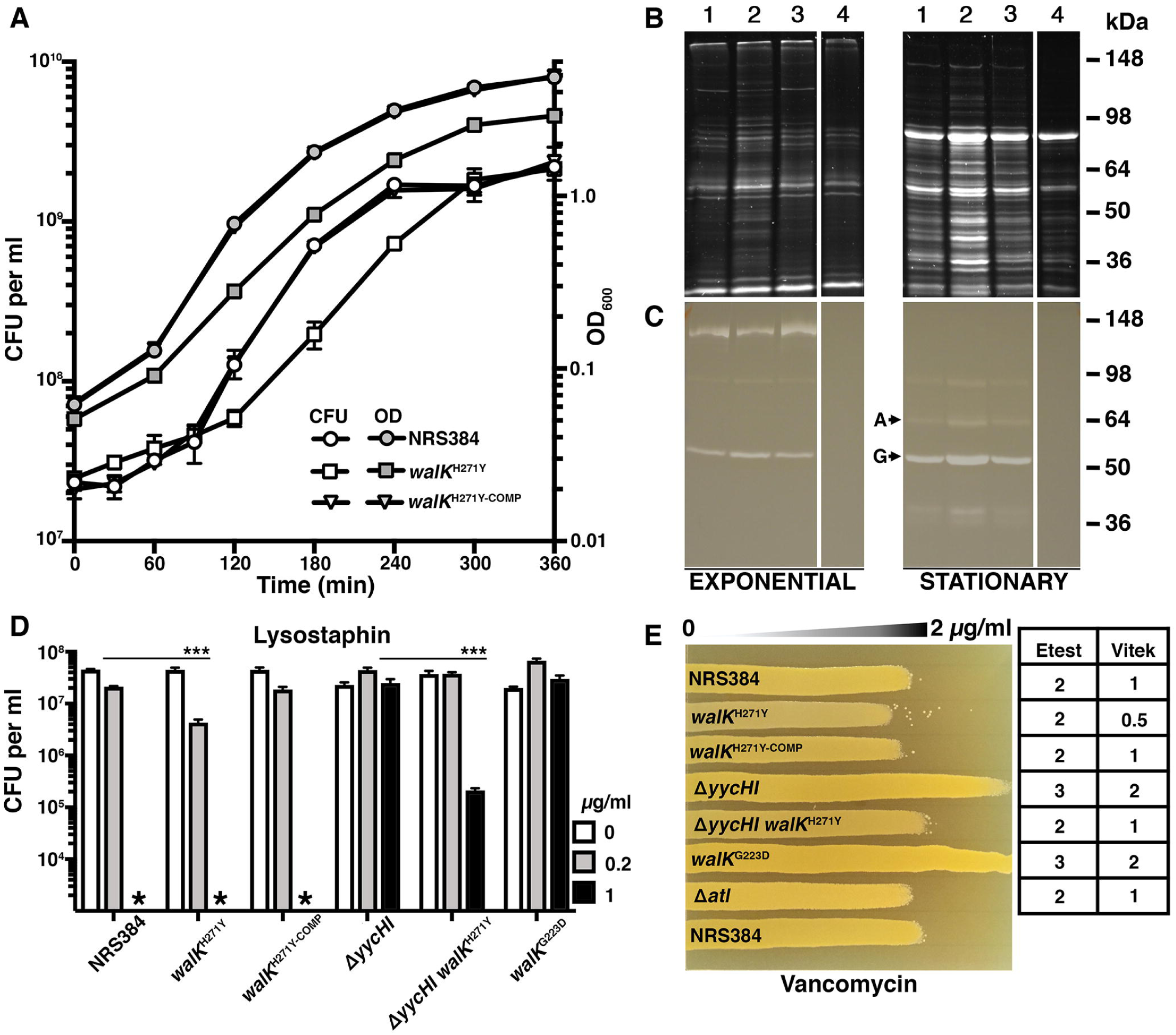
Phenotypic impact of WalK^H271Y^ mutation on *S. aureus.* **(A)** Growth kinetics of the WalK^H271Y^ mutant compared to the WT and complemented strains. Overnight cultures were diluted 1:100 into fresh TSB and grown at 37°C (200 rpm). The cultures were sampled at indicated time points for OD_600_ and CFU readings. The WalK^H271Y^strain exhibited altered growth kinetics compared to the wild-type and complemented strains, with the loss of a lag phase and a subsequent reduction in doubling time during exponential growth. Error bars indicate standard deviation (SD) from 3 biological replicates. **(B)** Analysis of the total exoproteome and Atl secretion. Supernatant proteins from exponential or stationary phase cultures were run on 10% SDS PAGE gels for SYPRO Ruby staining or **(C)** zymogram analysis through the incorporation of 1.5% *Micrococcus luteus* cells. The lane contents are: 1. NRS384, 2. WalK^H271Y^, 3. WalK^H271Y-COMP^ and 4. Δ *atl,* with ‘A’and ‘G’ representing amidase and glucosaminidase, respectively. The protein profile of an *atl* mutant was included as a control for Atl activity. **(D)** Impact of the mutations on the sensitivity to lysostaphin. An overnight culture of the indicated strains was diluted 1:100 into fresh TSB containing the indicated concentration of lysostaphin, sampled for CFU (Time 0) and incubated statically for 90 min at 37°C. Asterisk denotes below the limit of detection (10^3^ CFU/ml). Depicted are the mean ± SD of at least triplicate biological experiments. The null hypothesis (no difference in mean lysostaphin sensitivity between wild-type and *walK*^*H271Y*^ at 0.2μg/ml or Δ *yycHI* and Δ *yycHI wall<*^*H271Y*^ at 1 μg/ml) was rejected for P<0.05*** (Student’ s t-test). **(E)** Impact on vancomycin sensitivity. Agar plates were made with a 0 to 2μg/ml concentration gradient of vancomycin. Independent Etest and Vitek vancomycin MIC measurements are indicated next to the gradient plate.

### The exoproteome and Atl activity is altered in the WalK^H271Y^ mutant strain

Atl is a major peptidoglycan hydrolase produced by *S*. *aureus* that is positively controlled by WalKR ^16,26^. It is involved in daughter cell separation and also plays roles in primary attachment during biofilm formation and in the secretion of moonlighting proteins ^33^. Here, we analysed the secretion of proteins during exponential and stationary phase growth from the wild-type and the mutant WalK^H271Y^, WalK^H271Y-COMP^ and Δ *atl* strains. We observed an increase in protein abundance in the WalK^H271Y^mutant in both exponential and stationary phases, by comparison to the wild-type levels observed in the walKH271Y-COMP strain. In contrast, secreted protein levels were reduced in the Δ*atl*strain compared to the wild-type (Fig. 2b). We then directly assessed Atl exoproteome by zymogram analysis, using Micrococcus luteus cells as a substrate. The Δ*atl* strain activity in the displayed no visible lytic activity in contrast to the other strains, indicating that the majority of lysis is attributable to Atl^34^ (Fig. 2c). In stationary phase, there was an apparent increase in the amidase (63 kDa) and glucosaminidase (53 kDa) activity in samples from the *walK*^*H271Y*^ mutant. These data indicate that the *WalK*^H271Y^ mutation results in increased production and/or activity of Atl.

### Increased lysostaphin and vancomycin sensitivity of the WalK^H271Y^ mutant

WalKR regulates autolysis and directly influences sensitivity to vancomycin and lysostaphin ^16,22,23,32^.Here, we used lysostaphin and vancomycin sensitivity as indirect measures of WalKR activity. After a 90 min exposure to 0.2 μg/mL of lysostaphin, NRS384 exhibited a 0.5-logio_10_ reduction in cell viability, while the WalK^H271Y^ mutant showed a further 0.5-logio_10_ increase in sensitivity. Complementation returned lysostaphin sensitivity to wild-type levels (Fig. 2d). In contrast, the Δ*yycHI* mutant displayed increased lysostaphin resistance compared to the wild-type, consistent with YycHI positively regulating WalKR. Therefore, the knockout of *yycHI* leads to reduced WalKR-controlled autolytic activity (Fig. 2d) ^28^. The compensatory WalK^H271Y^ mutation in the Δ*yycHI* background did not fully restore lysostaphin sensitivity to wild-type levels. However, increased sensitivity to lysostaphin was observed at a higher concentration (1 μg/mL) with the Δ*yycHI walk*^H271Y^ strain 2- log_10_ less viable than the Δ*yycHI* strain (Fig. 2d) with the wild-type below the limit of detection. Building on this framework, we then analysed a *walK*^*G*^223^*D*^ mutant, which was previously shown to have reduced autophosphorylation/phosphotransfer between WalK and WalR ^22,23^. Here, this mutant showed increased resistance to lysostaphin, similar to that observed for the Δ*yycHI* mutant (Fig. 2d). We then analysed the impact of the mutations on vancomycin resistance, using antibiotic gradient plates (Fig. 2e). The wild-type, WalK^H271Y-COMP^ and Δ*atl* strains all showed similar levels of resistance. However, the *AyycHI* and WalK mutant strains showed increased resistance to vancomycin, while the WalK^H271Y^ strain exhibited increased sensitivity compared to the wild-type. Introduction of WalK^H271Y^in the Δ*yycHI* background restored sensitivity to wild-type levels (Fig. 2e). These findings were consistent with Vitek 2 and Etest analyses of the strains (Fig. 2e). Collectively, these results indicate that the WalK^H271Y^ mutation activates the WalKR system, resulting in increased sensitivity to lysostaphin and vancomycin.

### Structure of the WalK PAS domain

To gain further insight into the impact of the WalK^H271Y^ mutation, we determined a high-resolution structure of the cytoplasmic PAS domain. Domain boundaries for WalK-PAS were defined, based on limited proteolysis, as valine 251 to arginine 376. This PAS domain sequence (WalK-PAS^FULL^) was cloned and heterologously expressed as a fusion protein with an N-terminal glutathione S-transferase (GST) tag and a thrombin cleavage site. WalK- PAS^FULL^ was purified by affinity chromatography, then the affinity tag was removed prior to crystallisation of the PAS domain (Suppl. Fig. 1). A selenomethionine derivative was also generated to aid in phasing the diffraction data (Suppl. Fig. 2). The WalK-PAS^FULL^ structure was solved to 2.0Å, although a lack of density for the N- and C- termini precluded modelling residues 251-262 and 370-376, respectively. Density was also absent for residues 335-338, which represents a disordered loop in WalK-PAS^FULL^. Details of the diffraction data and structure statistics are summarised in Table SI.

The WalK-PAS^FULL^ structure has a typical PAS domain fold ^35^ comprising five antiparallel β-strands and four α-helices, with connecting loops between the helices and β-strands (Fig. 3a,b). The N-terminal region of the structure is composed of two short β-strands connected by a small loop. The remainder of the N-terminal region is comprised of four short helices connected by two large loops. The C-terminal portion of the PAS domain predominantly consists of a short β-strand followed by two larger antiparallel β-strands connected by a loop. Analysis of the surface electrostatic potential showed an uneven charge distribution on the WalK-PAS^FULL^ surface (Fig. 3c). In contrast to other PAS domains, a well-defined and continuous electron density was observed for a single zinc atom coordinated by a hitherto unknown metal-binding site (Fig. 3d). Notably, the Zn^2+^-binding site resides on the surface of the WalK-PAS^FULL^ domain, with access for Zn^2+^ from the surrounding solvent. The metal binding site comprises a single Zn^2+^ ion bound by the atoms Nδ1 from His271, 0δ1 from Asp274, Nδ1 from His364, and Oε2 from Glu368 in a slightly distorted tetrahedral coordination geometry (Fig. 3d, Suppl. Fig. 3a). The observed bond lengths are 2.20Å, 2.23Å, 2.23Å, and 2.24Å, respectively, and are typical for protein coordination of a Zn^2+^ ion ^36^. The Zn^2+^-binding site, along with the rest of the exposed region, exhibits a negatively-charged surface, while the remainder of the WalK-PAS^FULL^ surface is relatively neutral. Although the N-terminal region is well ordered in the structure, the mobility of this region may increase in the absence of the Zn^2+^ ion and the stabilising interactions conferred by His271 or Asp274.

**Figure 3.**
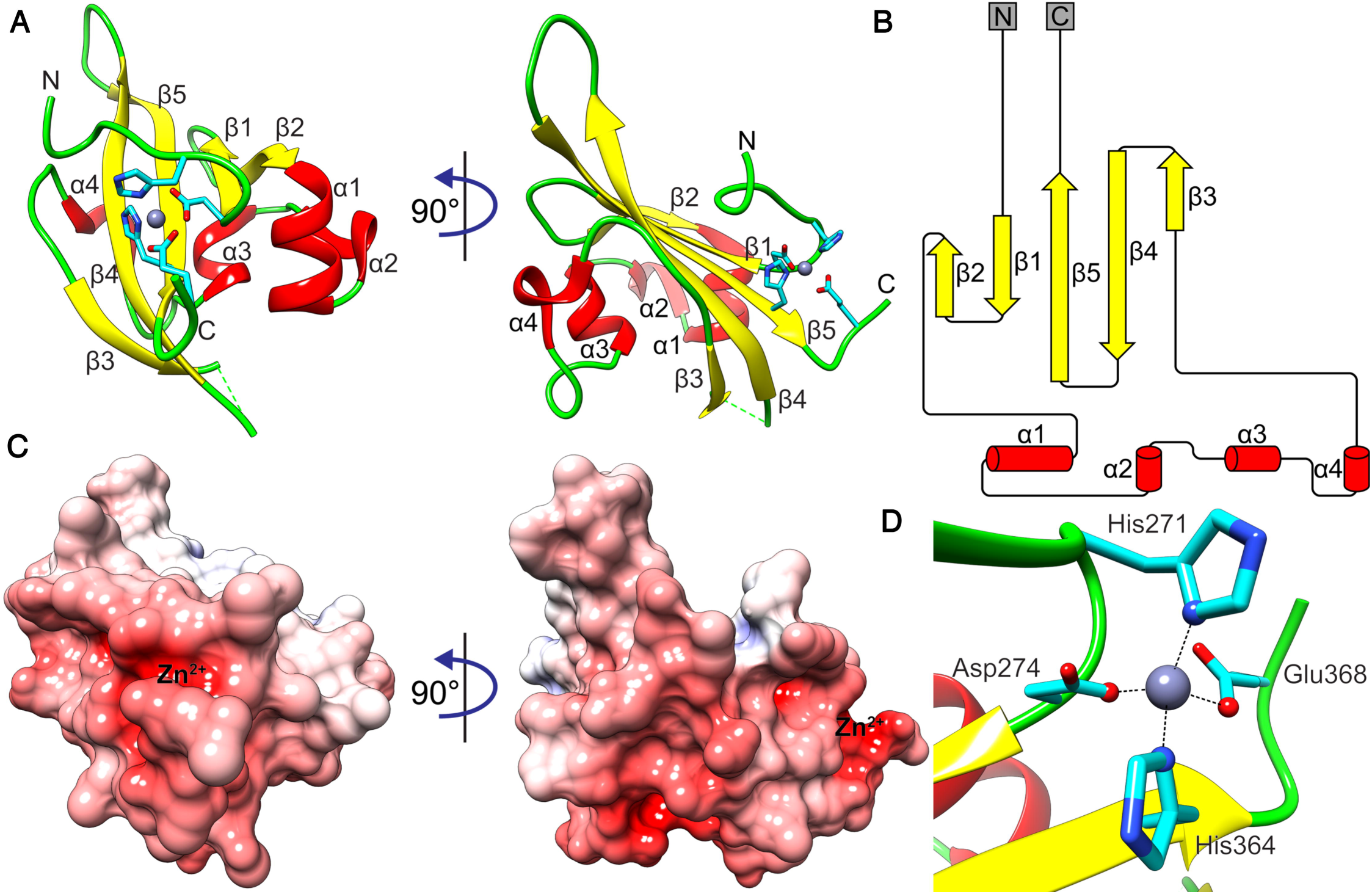
Crystal structure of *S. aureus* WalK-PAS^FULL^. **(A)** Cartoon representation of the crystal structure of WalK-PAS^FULL^. The α-helices are coloured red, β-strands yellow, and loops green. The bound Zn^2+^ is shown as a grey sphere and its coordinating residues as cyan sticks. The N- and C- termini of the structure are labeled. **(B)** Sequence and crystal structure-based secondary structure of WalK-PAS^FULL^ generated by Pro-origami^60^. α-Helices are shown as red cylinders and β-strands as yellow arrows. **(C)** Surface electrostatic potential of WalK-PAS^FULL^ shown in the same orientation as in (A). Positive and negative potentials are shown in blue and red, respectively, coloured continuously between –10 and 10 kT/e. Surface electrostatic potential was calculated using APBS ^61^; the calculation included the Zn^2+^ ion. **(D)** The Zn^2+^-binding site of WalK-PAS^FULL^. Metal-coordinating residues are shown as cyan sticks, with the atoms contributing to the interactions as spheres. The coordinating bonds are illustrated with black dashed lines.

### Analysis of WalK-PAS domain interaction with Zn^2+^

Further examination of the interaction of the PAS domain with Zn^2+^was performed using a truncation mutant (WalK-PAS^TRUNC^) in which the unstructured regions (15 residues deleted from the N-terminus and 6 residues from the C-terminus) were removed. The high-resolution structure of WalK-PAS^TRUNC^ was solved to 2.1 Å (Table S2). Comparison of the WalK-PAS^FULL^ and WalK-PAS^TRUNC^ structures revealed nearly identical PAS domain folds (Suppl. Fig. 3b,c), with a calculated RMSD of 0.48Å.

To investigate the Zn^2+^ binding capability of WalK-PAS, we conducted Zn^2+^ binding assays with both WalK-PAS^TRUNC^ and WalK-PAS^TRUNC H^271^Y^, followed by direct metal quantitation by inductively coupled plasma-mass spectrometry (ICP-MS). Zinc binding analyses of WalK-PAS^TRUNC^ revealed a metakprotein molar ratio of ˜0.3:1, suggesting that WalK-PAS was ˜30% Zn^2+^-bound at any given time. By contrast, no Zn^2+^ binding was observed for WalK-PAS^TRUNCH^271^Y^. Taken together, these data indicate that the H271Y mutation abolishes Zn^2+^ binding by the WalK PAS domain, supporting the inference that metal binding at this site influences WalK activity. Isothermal titration calorimetry (ITC) revealed that WalK-PAS^TRUNC^ binds a single Zn^2+^ ion with *K*_*d*_ ˜340 nM, indicating a moderate affinity interaction, consistent the ICP-MS analysis. Analysis of WalK-PAS^TRUNC H^271^Y^ by ITC did not detect Zn^2+^ binding, consistent with the ICP-MS result. These data suggest that Zn^2+^ is unlikely to serve a structural role in the WalK-PAS domain, since structural Zn^2+^ sites in proteins are typically formed by coordination spheres comprising Cys_4_ or Cys_2_-His-Cys ligands and they are associated with much higher binding affinities for Zn^2+^ (100 nM – 100 pM)^36^. By contrast, Glu and Asp residues are infrequently found in structural Zn^2+^ sites, with a recent analysis of NCBI Protein Data Bank noting a prevalence of ≤ 6.0% for relevant protein structures ^36^. Collectively, these data, in combination with the phenotypic observations for the *walk* ^H*271Y*^ mutant, strongly suggest that the metal binding site in WalK-PAS serves a regulatory role than a structural function.

### PAS-domain Zn^2+^-binding regulates WalK autophosphorylation

To elucidate the regulatory role of metal binding in the WalK-PAS domain, we examined the impact of the H271Y mutation on WalK autophosphorylation activity. Measurements were performed using the cytoplasmic domains of WalK^208608^ (WalK^CYT^), which has previously been studied using autophosphorylation assays ^10,23^. Here, we observed that the H271Y mutation (WalK^CYTH^271^Y^) resulted in a ˜50% increase in autophosphorylation, by comparison to WalK^CYT^ (Fig. 4a). Hence, these data indicate that Zn^2+^- binding by the PAS domain negatively regulates the autophosphorylation activity of WalK in *vitro,* consistent with the altered WalK activity observed by *in vivo* phenotypic analyses.

**Figure 4.**
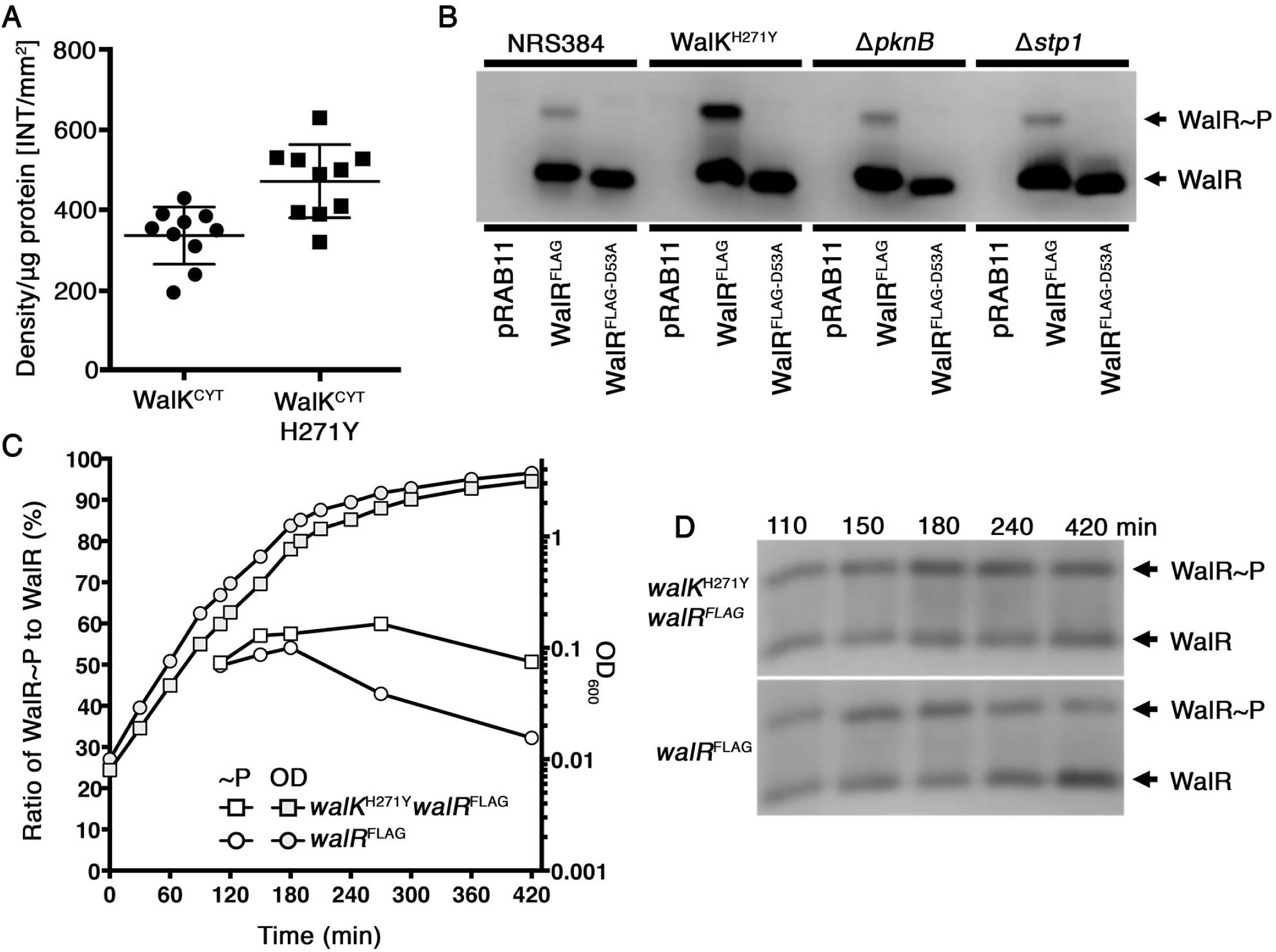
*In vitro* analysis of WalK autophosphorylation and *in vivo* analysis of WalR phosphorylation. **(A)** Autophosphorylation of WalK. Incubation of WalK^CYT^ and WalK^CYT^-^H271Y^ with [γ-^32^P]-ATP for 60 min shows H271Y mutation increases WalK autophosphorylation (p=0.0016**). Proteins were electrophoresed on 12% SDS-PAGE gels and the level of radiolabel incorporation quantified using autoradiography of the phosphorylated protein bands. Error bars SD. Null hypothesis (no difference between means) was rejected for p<0.01 (two-tailed, Mann-Whitney U- test). **(B)** Establishing phos-tag acrylamide for the analysis of WalR phosphorylation. Either pRABll (empty vector), WalR^FLAG^ (wild-type WalR with 3xFLAG tag), or WalR^D^53^A^-^FLAG^ (mutation to abolish D53 phosphorylation in WalR^FLAG^) were transformed into NRS384, WalK^H271Y^, Δ ***pknB*** or Δ ***stpl.*** Cultures were diluted into fresh TSB and grown to early stationary phase in the presence of 10 Hg/ml chloramphenicol and 0.4 μM ATc. Cells were processed as described in the Materials and Methods, with equivalent concentrations of protein loaded onto 8% SDS-PAGE gels containing Mn^2+^ Phos-tag (50 μM). WalR was detected by western blot with anti-FLAG M2 monoclonal antibody. **(C)** Analysis of chromosomally FLAG tagged WalR by phos-tag. The *walR* gene in either the wild-type or WalK^H271Y^ background was tagged with a 3x FLAG tag on the C-terminus. A growth profile (grey circle/square) and the phosphorylation status (white circle/square) of WalR was examined at time points equating to early, mid, late log and early stationary phase. The ratio of phosphorylated to non-phosphorylated WalR presented in (C) was determined by densitometry of phos-tag electrophoresised / western blotted samples shown in **(D).**

To complement the *in vitro* findings, we investigated the *in vivo* phosphorylation status of WalR. We first established the detection efficiency for the liable aspartic acid phosphorylation on residue D53 of WalR. This was achieved by constructing two WalR variants into the tetracycline inducible plasmid pRABll, containing either: (i) native WalR with a 3xFLAG tag on the C-terminus (WalR- FLAG); or (ii) a mutant variant of WalR-FLAG, wherein mutation of D53 to an alanine (D53A). The D53A mutation abolishes the potential for phosphorylation at this site. The above plasmids along with empty pRABll were transformed into the 5. *aureus* NRS384 wild-type, WalK^H271Y^, *Δstpl* (serine threonine phosphatase) and Δ*pknB* (serine threonine kinase). The latter two strains were included to differentiate the site of phosphorylation. PknB has been shown to phosphorylate WalR at residue T101^37^ while in *B. subtilis,* the deletion of Stp1 led to increased WalR phosphorylation at the site equivalent to WalR T101^38^. Here, we observed that D53 phosphorylation on WalR was absent in the WalR^D^53^ FLAG^ (Fig. 4b). Deletion of Δ*PknB* or Stpl did not alter D53 phosphorylation, indicating that WalK was the dominant contributor. Further, our analyses did not detect a second site of phosphorylation under these conditions, suggesting that T101 was not being phosphorylated or detected under the conditions tested. Increased phosphorylation of WalR was observed in the WalK^H271Y^ background (Fig. 4b), consistent with the autophosphorylation results (Fig. 4a). Building on this framework, we examined D53 phosphorylation in the native context using chromosomally-tagged strains (WaIR) in either wild-type or WalK *S*. *aureus.* The ability to modify the chromosomal copy of *walR* showed that the FLAG tag did not dramatically alter WalR activity. When analysed throughout growth, there was a striking increase in the phosphorylation of WalR in the WalK^H271Y^ strain over the wild-type from mid-log phase onwards, with the difference exaggerated as growth progressed into late log/early stationary phase (Fig. 4c,d). These findings are consistent with the WalK autophosphorylation assay (Fig. 4a) and show that the negative regulation of WalKR imposed by Zn^2+^ binding to the cytoplasmic PAS domain is important in dampening the response of WalK as the cells transition out of active growth.

### Conservation of metal binding sites in WalK is restricted within the Firmicutes

We next examined the potential conservation of the novel WalK PAS^CYT^ domain metal binding site across low G+C Gram-positive bacteria. Alignment of a selection of WalK proteins from different genera with the *S*. *aureus* WalK reference sequence revealed that H271 is conserved among the coagulase positive/coagulase negative staphylococci and enterococci, but not in streptococci or listeria, where it is replaced by a tyrosine residue (Fig. 5a). A Y271 residue was also present in two out of the five bacilli examined, including the well-studied strain, *B. subtilis* 168. Notably, the three additional metal coordinating residues were only conserved among staphylococci and enterococci, with all other genera having at least one deviation from the *S*. *aureus* consensus. These results suggest that conservation of metal binding by WalK in staphylococci and enterococci might enable additional regulatory control of the essential WalKR TCS. We next examined the structural alignment of the WalK-PAS^CYT^ domains of S. *aureus* with that from *Streptococcus mutans* (WalK ^SM^); the latter natively encodes the Y271 substitution and was crystallized in the context of the complete cytoplasmic domain ^39^ (Fig. 5b). Although the two PAS^CYT^ domains align with a relatively large RMSD of 2.31 Å, indicating significant structural differences, the largest deviations occur in the regions comprising the Zn^2+^-binding site of thes. *aureus* PAS^CYT^ domain. In *S*. *mutans* the WalK^CYTSM^ domain forms a leucine zipper dimeric interface with an adjacent monomer. Metal binding by *S*. *aureus* WalK-PAS^CYT^ is predicted to preclude formation of such an interface. The Zn^2+^ atom would be positioned near the center of the structure, resulting in steric clashes at the dimeric PAS^CYT^ domain interface that would likely impede WalK dimer interactions (Fig. 5c).

**Figure 5.**
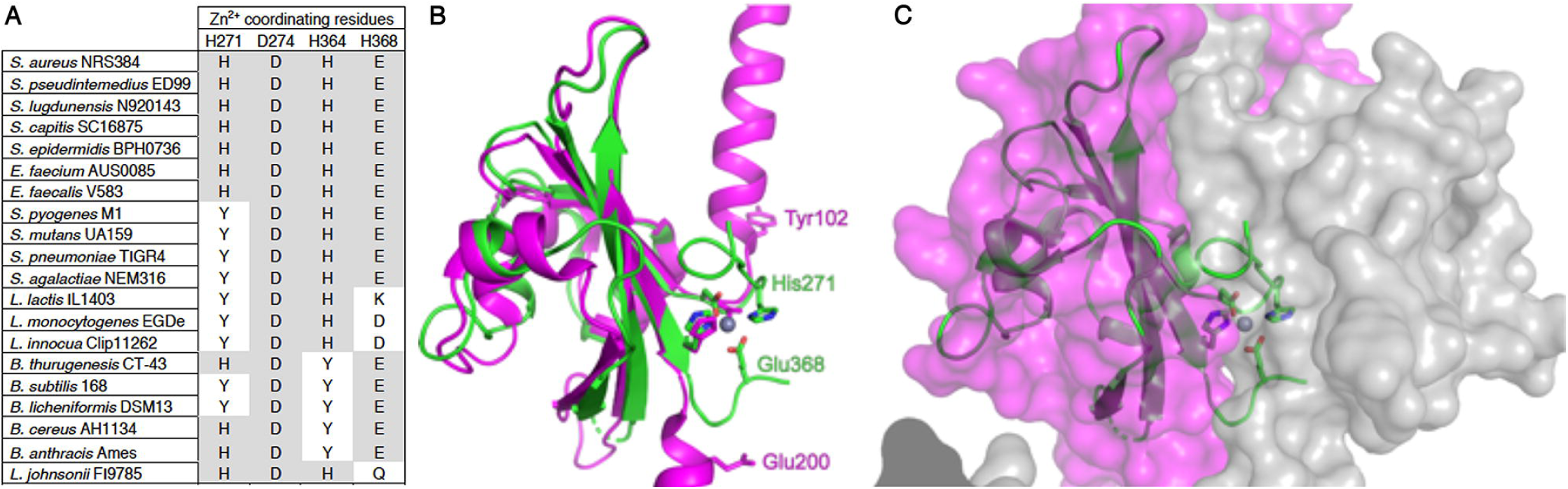
Structural comparison of WalK. **(A)** Comparison of metal coordinating residues from a range of Gram-positive bacteria with WalK orthologs. Protein sequences were aligned with ClustalW using default parameters. Residues which match the S. *aureus* consensus are highlighted in red. **(B)** Superposition of the crystal structure of WalK-PAS^FULL^ (green) with VicK (magenta, PDB: 4I5S) in cartoon representation. The Z^2+^-coordinating residues in WalK-PAS^FULL^ are shown as sticks, with the homologous residues in VicK shown in magenta. The bound Zn^2+^ ion is shown as a sphere. **(C)** Superposition of WalK-PAS^FULL^ (green) with the VicK_2_ homodimer shown by transparent surface representation (magenta/grey, PDB: 4I5S). The crystal structure of WalK-PAS^FULL^ is shown by cartoon representation with the Zn^2+^-binding site metal-coordinating residues shown as sticks and the bound ion as a sphere.

### Metal-induced conformational changes in WalK

To further understand the structural consequences of WalK Zn^2+^-binding and impact on kinase activity, we employed molecular dynamics (MD). MD simulations indicated that Zn^2+^-binding directly influences the relative positioning of the PAS and catalytic (CAT) domains. In the absence of Zn^2+^, the dihedral angle between the PAS^CYT^ and CAT domains in each monomer was ˜136° when viewed down the central axis of the kinase (measured as the dihedral angle between Cα atoms of residues 288, 271, 369 and 569, averaged between the two chains), while the average distance between the upper and a and averaged between the two chains) was 21.6 Å (Fig. 6a). In the presence of Zn^2+^, the relative dihedral angle increased to 175° while the intra-helical distance decreased to an average of 12.3 Å (Fig. 6b). The binding of the metal ion also stabilizes the PAS domain fold by bringing the N- and C-terminal regions into closer proximity. This is particularly evident for the N-terminal H271 (Tyr in WalK-PAS^SM^) and the C-terminal Glu368 residues. Collectively, our structural analyses suggest that metal binding in the PAS domain results in significant conformational changes. These metal-induced structural rearrangements provide a mechanistic basis for how conformational changes arise in WalK to negatively regulate its autokinase activity and subsequent signal transduction to the response regulator, WaIR.

**Figure 6.**
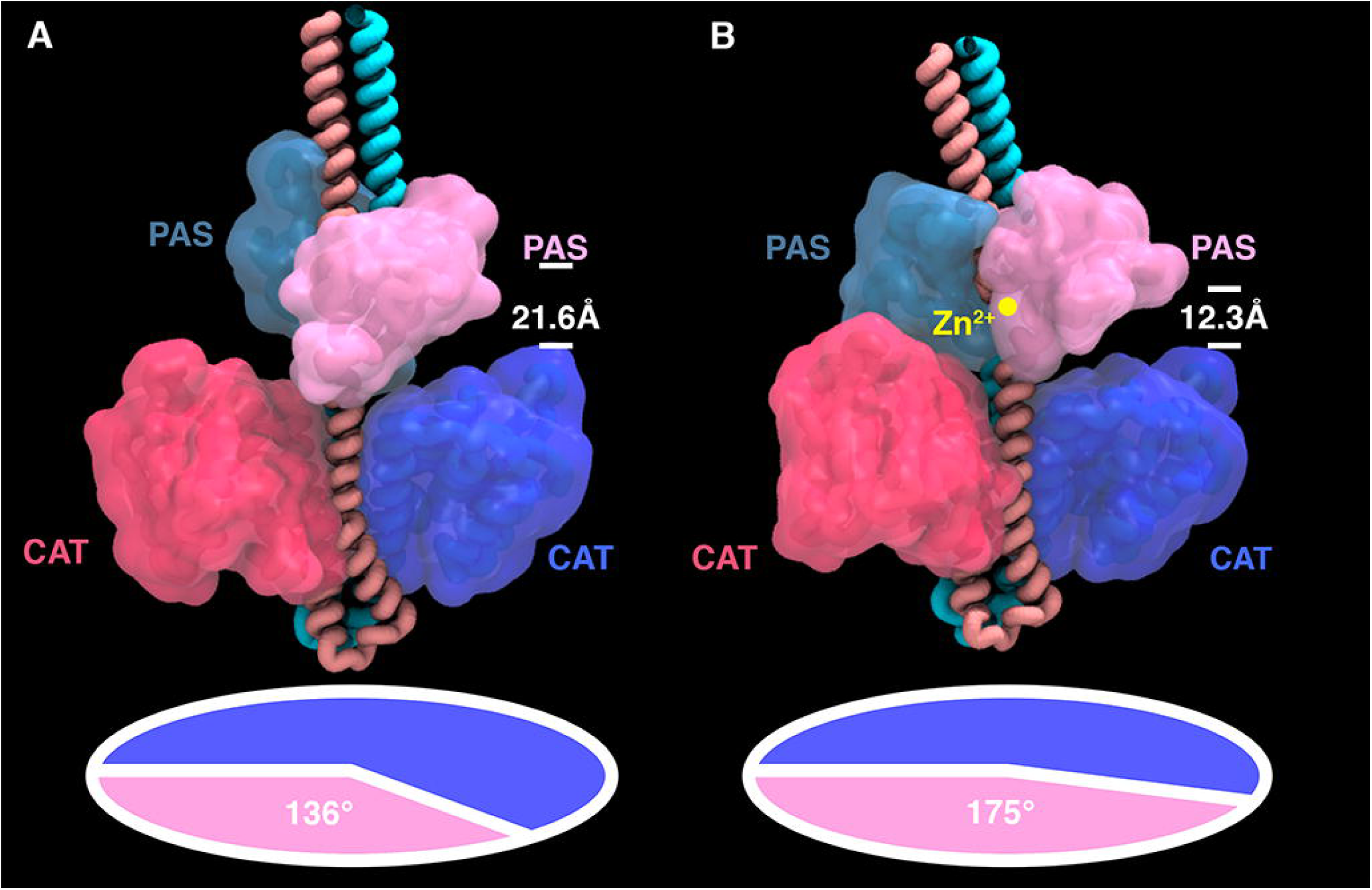
Molecular modelling of WalK in the presence or absence of Zn^2+^. Still images taken from molecular dynamics simulation of full-length, membrane-bound *S*. *aureus* WalK in the **(A)** absence and **(B)** presence of Zn^2+^, showing the predicted conformational changes induced in the WalK PAS and CAT domain upon metal binding.

## DISCUSSION

In this study, we provide the first evidence of a specific ligand for the WalKR system, opening new avenues to understand the function and essentiality of this regulon. The S. *aureus* WalKR two-component system was identified in the late 1990s, but the ligand(s) sensed by this histidine kinase and subsequent mechanisms of activation have remained elusive. One proposed model for WalK sensing has been through the recognition of the D-Ala-D-Ala moiety of Lipid II via the extra-cytoplasmic PAS domain ^6^. This molecule would be abundant at the site of septation during exponential growth, but become limiting upon the cessation of cellular replication. Although this scenario provides a link between activation of autolysin production, via phosphorylation of WalR, with division septum localisation of WalK, there remains a paucity of experimental evidence to support this model ^40^. Here, we have shown that the presence of a metal binding site within the PAS^CYT^ domain of WalK directly influences the activation status of the protein. Abrogation of *in vitro* Zn^2+^-binding capacity of WalK increased the autophosphorylation of WalK and *in vivo* phosphotransfer to WalR. There are several key examples of regulation of histidine kinase activity via PAS domain ligand-binding, such as oxygen sensing by *Bradyrhizobium japonicum* FixL, wherein heme binding by the cytoplasmic PAS domain regulates nitrogen fixation under reduced oxygen tensions ^35^; oxygen sensing by *Staphylococcus carnosus* NreB, in which the PAS^CYT^ domain contains an oxygen-labile iron-sulfur cluster ^41^; and redox status sensing by *Azotobacter vinelandii* NifL, where the PAS^CYT^ domain binds nicotinamide adenine dinucleotide ^42^. Direct metal ion binding has previously been observed in extracellular PAS domains, such as PhoQ. from *Salmonella typhimurium,* which senses the cations Ca^2+^ and Mg^2^+^43^. However, to the best of our knowledge, metal binding by a cytoplasmic PAS domain has not previously been reported and appears to be a highly restricted attribute among the staphylococci and enterococci, despite WalK being conserved among the Firmicutes. For instance, the WalK PAS^CYT^ domain in *Streptococcus mutans* has a naturally occurring tyrosine at residue 271 and structural analysis shows no evidence of metal binding^39^.

Recapitulation of this mutation *in vivo* resulted in phenotypes associated with activation of WalK (e.g. sensitivity to lysostaphin and vancomycin, increased hemolysis and activity/production of the major autolysin Atl) along with the loss of lag phase upon inoculation into fresh media. This latter phenotype would be consistent with the requirement for accumulation of a ligand sensed by the extracellular PAS domain leading to the activation of WalKR, and subsequent autolysin production. By contrast with *B. subtilis,* YycHI are activators of the *S*. *aureus* WalK system ^28,29^. Consequently, the metal-dependent regulation of the WalK-PAS domain in *S*. *aureus* may serve as a dynamic constraint on kinase activity. Additional levels of regulation of the WalKR operon have also been identified. These include a second site of phosphorylation, residue T101 on WalR, by the serine threonine kinase known as Stkl or PknB ^37^, and the interaction of SpdC (previously called lysostaphin resistance factor A – *lyrA)* ^44^ with WalK, which negatively regulate genes under the control of WalR^45^. Both PknB and SpdC are localised to the division septum similar to WalK ^31,37^, highlighting the complexity of this regulatory axis. The establishment of phos-tag acrylamide to analyse the phosphorylation status of WalR *in vivo* is a powerful tool to investigate the regulation of this essential system (Fig. 4).

The PAS^CYT^ domain of *S*. *aureus* WalK binds Zn^2+^ with only moderate affinity, suggesting a regulatory role for metal binding rather than an obligate structural function. In this manner, it is possible that this site only has transient interactions with Zn^2+^ thereby facilitating a continuum of WalK activation states rather than serving as a binary switch. Disruption of PAS^CYT^ domain dimerization upon metal binding could impact WalK activity in a similar manner to the L100R mutation in *Streptococcus pneumoniae* ^46^. The L100R mutation destabilizes *S*. *pneumoniae* PAS dimerization, leading to a loss of autophosphorylation activity. Our structural analyses suggest that Zn^2+^ binding induces large conformational changes that alter the relative positions of the PAS^cyT^ and catalytic domains are induced by Zn^2+^-binding. These observations provide a plausible mechanistic basis for the reduction in WalK function, although the precise mechanism remains to be elucidated.

Intriguingly, although the amino acid sequence differences between WalK from the staphylococci and enterococci compared to other Firmicutes are small, these differences appear to have major functional consequences. This may suggest differing or additional regulatory roles for WalK beyond peptidoglycan biosynthesis in these genera. Potential regulatory roles for WalK could include contributing to maintenance of intracellular metal homeostasis. Sensing of intracellular Zn^2+^ abundance by WalK, could influence the import and/or efflux of cytoplasmic Zn^2+^, via WalR mediated phosphorylation. However, transcriptome studies of *S*. *aureus* WalKR mutants commonly highlight genes and pathways involved in nucleotide metabolism, and not those associated with metal ion transport ^22,26^. Nonetheless, indirect metal-dependent regulation might influence such pathways. One potential mechanism would be via Stp1, a Mn^2+^-dependent phosphatase, and its cognate kinase PknB, which interacts with WalKR and influences its activity. Further insight into the role of metal coordination in regulating this histidine kinase is necessary to understand the essentially of the system in *S*. *aureus.*

## METHODS

### Strains, primers, reagents and media

Bacterial strains/plasmids and primers (IDT) used are described in Table S2 and Table S3, respectively. *S*. *aureus* were routinely grown on Brain Heart Infusion (BHI) agar (Difco) or in Tryptone Soy Broth (TSB) (Oxiod) at 37°C with shaking at 200 rpm. For the selection of pIMAY-Z containing strains, BHI agar was supplemented with 10 μg/ml chloramphenicol and 100 μg/ml X-gal (BHIA-CX). For protein expression, Terrific Broth (TB) was used (10 g/L tryptone, 24 g/L yeast extract, 10 g/L glucose, 0.17 M KH_2_P0_4_and 0.72 M K_2_HP0_4_). Restriction enzymes, Phusion DNA polymerase and T4 DNA ligase were purchased from New England Biolabs. Phire Hotstart II (for colony PCR) was purchased from Thermo Fisher. Genomic DNA from *S*. *aureus* was isolated from 1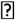ml of an overnight culture (DNeasy Blood and Tissue Kit—Qiagen) pretreated with 100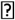μg of lysostaphin (Sigma cat. no. L7386). Lysostaphin sensitivity assays were performed as described^32^.

### *S. aureus* site-directed mutagenesis by allelic exchange

The *walK*^*H271Y*^ mutation was recombined into the chromosomal copy of *walK* in the USA300 background (NRS384) by allelic exchange. The region encompassing *walK* was amplified by SOE-PCR to introduce a neutral change in the third nucleotide of codon 270 and the first nucleotide in codon 271 with primer set IM7/IM8/IM9/IM10 (Table S4). The resultant 2.7 kb product was recombined into pIMAY-Z by the seamless ligation cloning extract (SLiCE) method ^32,47^ and transformed into *E. coli* IM08B ^48^. The sequence confirmed construct was extracted from *E. coli,* ethanol precipitated and transformed into electrocompetent NRS384 ^49^. Allelic exchange was performed as described ^48^, with the mutation screened using primer pair IM11/IM12 with an annealing temperature of 65°C. Reversion of the *walK*^*H271Y*^ mutation to *walK* ^C^0^MP^-^Pstl^ was achieved through allelic exchange in NRS384 *walK* ^H271Y^. A marked *walK* allele was constructed through the introduction of a silent *Pst\* site (*walK* nucleotide 1302, A to G) into *walK* by SOE-PCR with primer set IM7/IM58/IM59/IM10. A 3xFLAG tag was introduced onto the C-terminus of WalR by SOE-PCR with primers IM31/IM108/IM109/IM40. Deletion of genes or insertion of a FLAG tag was performed by SOE-PCR with the product cloned into pIMAY-Z by SLiCE with the following primer sets: *Δatl* gene (IM96/IM97/IM98/IM99: entire gene), Δ*yycHI* (IM54/IM78/IM79/IM75: from codon 5 of *yycH* to the stop codon of *yycI), Δstpl* (IM251/IM252/IM253/IM254: from the start codon leaving the last 8 amino acids), Δ*pknB* (IM255/IM256/IM257/IM258: entire gene) or *walR*^FLAG^ (IM31/1M108/1M109/1M40). The *walK*^*2230*^ mutation was directly amplified by PCR on JKD6008 genomic DNA with IM7/IM10 (Table S4). The purified pIMAY-Z construct was transformed into NRS384 after passage through IM08B (Table S4). Allelic exchange was performed as described above. To screen for the *walk*^J:LAG^, IM111/IM181 was used to identify the insertion. For WalK^G^223^D^ after allelic exchange, white colonies were screened for decreased sensitivity to vancomycin. Genome sequencing and analysis of the isolates was conducted as described previously^32^.

### Construction of anhydrotetracycline (ATc) inducible WalR-FLAG

The WalR^FLAG^ gene was PCR amplified from NRS384 *walR*^FLAG^ genomic DNA with primers IM280/IM305 (wild-type) or by SOE-PCR with primers IM280/1M278/1M279/1M305 to introduce the *walR*^D^53^A^ mutation. The products were digested with KpnI/BamHI and ligated into Kpnl/Bglll digested pRAB11. Ligations were transformed into IM08B to produce either pRAB11 *walR* ^FLAG^ or Prab11 *walR* ^D^53^AFLAG^. Plasmids (pRAB11, pRAB11 *walR* ^FLAG^ or pRAB11 *walR* ^D^53^AFLAG^) were transformed into NRS384, NRS384 *walR* ^H271Y^, NRS384 *Δ stp1* or NRS384 *Δ pknB* and selected on BHIA- CX at 37°C.

### Antibiotic resistance profiling

For vancomycin gradient assays were performed as described except a 0 to 2 μg/ml gradient was used ^50^. After 24 h incubation at 37°C, plates were imaged. With the exception of Vitek 2 and Etests, all antibiotic susceptibility testing of strains was performed in triplicate. To access the susceptibility to a range of antibiotics, strains were tested on a Vitek 2 Gram Positive ID card (AST-P612; Biomerieux) as per manufacturer’s instructions. Extended glycopeptide susceptibilities were determined with vancomycin Etest strips (Biomerieux) using a 2.0 McFarland inoculum on thick BHI agar and incubation of 48 h at 37°C.

### Supernatant protein precipitation and SDS-PAGE analysis

Supernatant proteins from overnight culture or OD_600_ = 0.8 TSB cultures were precipitated with 10% trichloroacetic acid at 4°C for 1 h. Pellets were washed with 1 ml of ice-cold acetone and air dried at room temperature. Precipitated proteins were resuspended in 2x SDS-PAGE loading buffer and prior to loading the samples were equilibrated with an equal volume of 1 M Tris.Cl [8]. Zymogram analysis of supernatant proteins was conducted as described by Monk etal.^51^.

### Cloning, expression and purification of the GST-WalK PAS domain in *E. coli*

WalK-PAS^FULL^ (residues Val251-Arg376) was PCR amplified from NRS384 genomic DNA with oligonucleotides WalK-PAS^FULL^-F/WalK-PAS^FULL^-R and cloned into pGEX-2T (*BamH\/EcoR\)* to yield a N-terminally tagged GST-PAS construct. For WalK-PAS^TRUNC^and WalK-PAS^TRUNCH^271^Y^ (residues Asp266 to Glu371), codon-optimised genes were ordered from GeneArt, and these were sub-cloned as *BamH\/EcoR\* fragments into pGEX-2T yielding pGEX-2T(WalK-PAS^TRUNC^) and pGEX-2T(WalK-PAS^TRUNC H^271^Y^). Overnight cultures of BL21 containing the different GST constructs were diluted 1:100 into 2 L of TB at 37°C at 180 rpm and grown to OD_600_ of 0.8. The culture was then induced with 0.4 mM IPTG and shifted to 16°C at 120 rpm for 10 h (final OD_600_ of ˜3). The cells were harvested by centrifugation at 5,000 *g* for 15 min at 4°C. The pellet was resuspended in 50 ml of GST lysis buffer (50 mM Tris.Cl [8.0], 500 mM NaCl, 1 mM EDTA) containing 1 mg/ml lysozyme plus 1U/ml DNase A and incubated for 30 min on ice. Cells were lysed at 40 kpsi in a cell disruptor (Constant Systems), then cell debris was removed by centrifugation at 39,000 *g* for 30 min at 4°C. The supernatant was collected and passed through GST-affinity resin in a gravity-flow column (Bio-Rad). The resin was then washed with two column volumes of GST lysis buffer followed by equilibration with 1 column volume of TCB (20 mM Tris [8.0], 500 mM NaCl, 1 mM CaCl_2_). WalK-PAS was eluted by on-column digestion with 20 ml. of TCB containing 200 U of thrombin. Thrombin digestion was carried out by incubating the column at room temperature for 30 min followed by 30 min at 37°C and 60 min at room temperature. The liberated PAS domain was purified by size exclusion chromatography using a Superdex 75 column. Purified PAS domains were concentrated using an Amicon centrifugal concentrator with a 10-kDa size cut-off. Protein purity was always greater than 95% as determined by SDS-PAGE with a yield of about 20 mg/L.

### Expression and purification of selenomethionine-labelled WalK-PAS domain

A single colony of BL21 + pGEX-2T(WalK-PAS^FULL^) was picked and used to inoculate 10 mL of pre-warmed L-broth (LB) at 37°C for 6 h (180 rpm). Minimal media (100 mL) was inoculated with 500 μL of this pre-culture. Cells were grown overnight at 37°C with shaking at 180 rpm, then 25 mL of culture was added to 2 L minimal medium and incubated at 30°C. After further growth to OD_600_ ˜ 0.7, amino acid stock solution containing selenomethionine was added, then cells were grown for 1 h before induction with 0.4 mM IPTG. Cells were harvested by centrifugation at 6,000 *g* for 20 min at 4°C and the resulting cell pellet stored at – 80° C. The SeMet-labelled WalK-PAS^FULL^ was purified as described above. Labelling was confirmed by peptide mapping through matrix-assisted laser desorption/ionization (MALDI) mass spectrometry (MS). A total of 10 μ g of purified protein was subjected to digestion for 16 h with 125 ng porcine trypsin (MS-grade, Promega) in 200 mM tetraetyl ammonium bicarbonate [8.0] containing 10 % acetonitrile. The peptide digest was mixed in a 1:1 v/v using solution of α-cyano-4 hydroxycinamic and matrix (5 mg/ml in 50 % acetonitrile in 0.1 % trifluoroacetic acid), spotted directly onto stainless steel MALDI target and MALDI-time of flight (TOF)/TOF spectra acquired using a Model 4700 Proteomic Analyzer (Applied Biosystems). For the digested peptides, the mass spectrometer was operated in reflector positive ionization mode using a m/z range of 700–4000. The MS peak list was extracted in GPS explorer software using the default parameters. A list of theoretical tryptic peptides obtained with the program GPMAW (allowing for one missed cleavage) was used to interpret the MS spectra based on an average increase in m/z of 47 Da for each selenomethionine residue.

### Cloning, expression and purification of the cytoplasmic domains of His_6_-WalK in *E. coli*

The cytoplasmic region of WalK (residues 208-608 ^TAA^), with or without the H271Y mutation, was PCR amplified from NRS384 genomic DNA with primers WalK208-F/WalK^TAA^-R (WalK^CYT^) or WalK 208F/IM8/IM9/WalK^TAA^-R (WalK^CYTH^271^Y^), with the latter assembled by SOE-PCR. The products a After transfer of the sequence-verified construct into BL21, an overnight culture in TB of pET19b(WalK^CYT^) or pET19b(WalK^CYTH^271^Y^) was diluted 1:100 in 2 L of TB. Cells were grown at 37°C and 180 rpm until OD_600_ ˜ 0.2 and then the growth temperature was reduced to 16°C. Protein production was induced with 0.2 mM IPTG at an OD_600_ ≈1, then cultures were grown overnight at 16°C. Cells were harvested by centrifugation at 8000 *g* for 8 min at 4°C, then stored at −80°C. Frozen cells were resuspended in native lysis buffer (50 mM Tris [7.5], 500 mM NaCl, 25 mM imidazole, 5 mM EDTA), passed through a cell disruptor (Constant Systems) at 35 kpsi, then cell debris removed by centrifugation at 39,000 *g* for 30 min at 4°C. The supernatant was loaded on a Ni-NTA column equilibrated with equilibration buffer (25 mM Tris [7.5], 25 mM imidazole, 500 mM NaCl). The column was washed with equilibration buffer followed by a wash step with pre-elution buffer (25 mM Tris [7.5], 90 mM imidazole, 500 mM NaCl, 5 mM EDTA). WalK protein was then eluted in the same buffer containing 250 mM imidazole and immediately desalted over a HiPrep 26/10 desalting column into desalt buffer (25 mM Tris [7.5], 150 mM NaCl, 2% glycerol).

### Crystalisation of the WalK PAS domain

The purified WalK-PAS^FULL^was concentrated to 15 mg/ml. and then screened for optimal crystallisation conditions. The protein was crystallised from conditions using the PACT screen (Molecular Dimensions). Plates were set up using a Mosquito crystallization robot (TTP Labtech). Crystallisation conditions were then refined with the best crystals formed in PAS buffer (100 mM Tris.Cl [8.0], 20 mM ZnCl_2_, and 26 % (w/v) PEG 3350) at 20°C. Notably, the presence of ZnCl_2_ was essential for crystal formation. WalK-PAS^FULL^ crystals were mounted using a 15-μM-nylon cryo-loop (Hampton Research) and soaked in cryoprotectant solutions consisting of the crystallisation PAS buffer containing 20% PEG 400 for 10 min each and flash-frozen in liquid nitrogen and stored in liquid nitrogen for X-ray diffraction studies. An initial native X-ray data set for WalK-PAS^FULL^ was collected in a cryo-stream (100 K) using an R-AXIS IV^t+^ image plate detector and CuKα radiation from a Rigaku free rotating anode generator (Rigaku/MSC). A total of 360 images were collected for the full dataset at 1D oscillation and 10- min exposure at a distance of 90 mm from the crystal to detector. The raw data was autoindexed, integrated and scaled using Scala in the CCP4 software package. The shorter WalK-PAS^TRUNC^ domain (residues 266-371) was crystallised using the same conditions.

### Model building and refinement

The structure of the PAS domain was solved by single-wavelength anomalous dispersion (SAD) using SeMet-labelled WalK-PAS^FULL^. A single crystal was used to collect a highly redundant dataset at the peak wavelength for selenium at the Australian Synchrotron (Table S1). Data was processed with MOSFLM^52^ and scaled with SCALA^53^. Heavy atom sites were found and refined to find the phase, and an initial model built using the AutoSol program in PHENIX^54^. The resulting initial model was then subjected to multiple rounds of refinement in PHENIX and rebuilding in COOT^55^ using a higher-resolution dataset collected with longer exposure times. The structure of the unlabeled shorter version of the WalK-PAS^TRUNC^ domain was subsequently determined by molecular replacement using PHASER^52^. Phasing and refinement statistics are given in Table SI.

### Molecular dynamics

A membrane–bound homology model of the WalK complex (sequence based on Uniprot entry Q2G2U4) was constructed with VMD ^56^ and SwissModel (https://swissmodel.expasy.org) using PDB crystal structures 4MN6 (residues 266 to 320, identity 100%), 5IS1 (residues 33 to 182, identity 100%) and 4I5S (residues 223 to 599, identity 44%) as templates ^57^. Missing regions 1 to 33, 183 to 222 and 600 to 608 were independently modelled using secondary structure prediction program *Psipred* (http://bioinf.cs.ucl.ac.uk/psipred/) as a guide as no significant similar structures were available. The missing N terminal and middle sections were modelled as trans-membrane helices, while the missing C-terminus was included as disordered (Suppl. Fig. 4). Models were fully solvated, ionised with 0.15 M NaCl, and embedded through a phosphatidylcholine (POPC) lipid bilayer. One version had Zn^2+^ ions bound to each of the cytoplasmic PAS domains, while the other was metal free. Initial dimensions were 127 x 127 x 260 Åcontaining 385994 and 385992 atoms respectively. Molecular simulations were performed using NAMD2.12 ^58^ with the CHARM36 force field ^59^.

Simulations were run with periodic boundary conditions using the ensembles at 37°C and 1 barpressure employing Langevin dynamics. Long-range Coulomb forces were computed with the Particle Mesh Ewald method with a grid spacing of 1Å. Time steps of 2 fs were used with non-bonded interactions calculated every 2 fs and full electrostatics every 4 fs while hydrogens were constrained using the SHAKE algorithm. The cut-off distance was 12 Åwith a switching distance of 10 Åand a pair-list distance of 14Å. Pressure was controlled to 1 atm using the Nose-Hoover Langevin piston method employing a piston period of 100 fs and piston decay of 50 fs. Trajectory frames were captured every 100 ps. The zinc-free model was simulated for 160 ns while the Zn^2+^- bound model was simulated for 200 ns. Simulations were unconstrained apart from weak harmonic constraints holding the Zn^2+^ ions in the bound position in the zinc-containing model. Dihedral angles between the cytoplasmic PAS and catalytic domains were measured over the course of the simulations with VMD.

### MALLS analysis for molecular mass determination

PAS domains were analyzed using multi-angle laser light scattering (MALLS) to determine their mass and level of polydispersity. Proteins were first separated by gel filtration on a Superdex 75 column, then eluted protein were passed through a miniDAWN light scattering detector and OptiLab refractometer (Wyatt Technology). Weight average molecular masses were determined from the refractive index and light scattering data using the Debye fitting method (ASTRA software package, Wyatt technology).

### *In vitro* Zn^2+^-loading assays

Metal loading assays were performed on purified apo-WalK-PAS^TRUNC^ and WalK-PAS^TRUNC H271Y^ (30 μM) by mixing with 10-fold molar excess Zn^2+^ (300 *\iM* ZnS0_4_) in a total volume of 2 ml in 20 mM MOPS [7.2], 100 mM NaCl for 60 min at 4°C. The sample was desalted on a PD10 column (GE Healthcare) into the above buffer, and then the protein concentration determined. Samples containing 10 μM total protein were prepared in 3.5% HNO_3_ and boiled for 15 min at 95°C. Samples were then cooled and centrifuged for 20 min at 14,000 g. The supernatant was then analysed by ICP-MS (Agilent 8900 QQQ), and the protein/metal ratio was determined.

### Isothermal titration calorimetry

The interaction between WalK-PAS^TRUNC^ and WalK-PAS^TRUNC H^271^Y^ was analysed via ITC using a MicroCal iTC200 calorimeter (Malvern Panalytical). The cell contained 50 μM protein in ITC buffer (50 mM MES [6] and 300 mM NaCl) and the syringe contained 3 mM ZnCI_2_ in ITC buffer. The titration was performed at 25°C with 16 injections of 3 μl with a spacing of 2 min between injections. Titrations were repeated three times. ITC data were fitted using Origin. A single-site binding model was used to fit the data to obtain the stoichiometry (*n*), enthalpy *{ΔH),* and binding affinity (*K*_d_).

### Autophosphorylation assay

WalK^CYT^or WalK^CYT H^271^Y^ (1 μg) were incubated at room temperature in 15 μl phosphorylation buffer (25 mM Tris, 300 mM NaCl, 1 mM TCEP, 20 mM KCl, 10 mM MgCI_2_, pH 8). Phosphorylation reactions were started by adding 1 μl of radiolabelled ATP mixture (2.5 μCi [y-^32^P]-ATP and 5 μM ATP) to the protein sample, which was then incubated for 60 min at room temperature. Reactions were stopped by adding 5 μl of 3x SDS-loading buffer, then samples were analysed on a 12% SDS-PAGE gel, followed by autoradiography. The intensity of phosphorylated protein bands was determined using Quantity One software (Bio-Rad).

### Detection of WalR phosphorylation using Phos-tag SDS-PAGE and Western Blot

Overnight cultures of NRS384, WalK^H271Y^, Δ *stpl* or Δ *pknB* containing either the empty vector pRAB11, pRAB11 *WalK* ^FLAG^ or pRAB11 *WalK* ^D53A FLAG^ were diluted 1:100 into 100 ml of TSB containing 10 μ g/ml_ chloramphenicol and 0.4 μ M ATc. Cultures were then grown to the start of stationary phase (OD_600_ ˜4.0). For the chromosomally tagged WalR^FLAG^ strains, overnight TSB cultures were diluted to OD_6oo_ = 0.01 in 1 L of TSB and samples were taken after 110 (early log), 150 (mid log), 180 (mid log), 240 (late log) and 420 min (early stationary phase). Samples were mixed with one sample volume of ice cold ethanokacetone and harvested by centrifugation at 7,300 *g* for 5 min at 4°C. The cells were washed with 20 ml of milliQ water and resuspended in 500 μ L of TBS (50 mM Tris.CI [7.5], 150 mM NaCl). Cells were disrupted by bead beating three times at 5,000 rpm for 30 s (Precellys 24, Bertin Instruments) and then the lysates were centrifuged at 11,000 *g* for 5 min at 4°C. A total of 25 μ g of protein was loaded on an 8% SDS-PAGE gel containing 50 μ M Phos-tag acrylamide (Wako Chemicals) and 100 μ M MnCl_2_. The gel was run according to the manufacturer’s instructions (Wako Chemicals). To remove manganese ions after electrophoresis, the gel was washed two times for 15 min with transfer buffer (25 mM Tris [8.3], 192 mM glycine, 20% methanol) containing 1 mM EDTA and once with transfer buffer without EDTA. The separated proteins were blotted onto a PVDF membrane using the Trans-Blot® Turbo™ transfer system (Bio-Rad) according to the manufacturer’s instructions. The membrane was treated with blocking buffer (5% EasyBlocker (GeneTex) in TBS, 0.05% Tween 20) for 16 h at 4°C and then with blocking buffer containing 1:500 mouse anti-FLAG® M2-Peroxidase (HRP) monoclonal antibody (Sigma) for 1 h at room temperature. The membrane was washed three times with TBS containing 0.05% Tween 20 and bound antibody was detected using the WesternSure® PREMIUM Chemiluminescent Substrate and the C-DiGit® Blot Scanner (LI-COR Biotechnology). The ratio of phosphorylated WalR was calculated by quantification of the western blot bands using GelAnalyzer 2010a.

### Data availability

All sequencing data used in this study have been deposited in the National Center for Biotechnology Information BioProject database and are accessible through the BioProject accession number PRJNA486581. Atomic coordinates and data for the cytoplasmic PAS domain of WalK have been deposited in the Protein Data Bank under accession numbers 4MN5 (WalK-PAS^FULL^; residues 251-376) and 4MN6 (WalK-PAS^TRUNC^; residues 266–371).

## Acknowledgements

We acknowledge funding from the Australian National Health and Medical Research Council (Project Grant GNT1010776 and Principal Research Fellowship GNT1044414 to G.F.K.; Senior Research Fellowship GNT1136021 to B.M.C.; Project Grants GNT1049192, GNT1129589 and Senior Research Fellowship GNT1105525 to T.P.S.; and Practioner Research Fellowship GNT1105905 to B.P.H.).

## Supplementary Figures

**Suppl Fig 1. Purification of recombinant WalK-PAS^FULL^. (A)** SDS-PAGE gel showing different stages in the purification of WalK-PAS^FULL^. Lane 1: whole cell lysate; lanes 2 and 3: soluble and insoluble fractions, respectively, post-cell disruption; 4: flow-through from loading soluble fraction onto GST-agarose column; 5: wash of GST-agarose column; 6: GST-agarose beads with bound GST- WalK-PAS; 7: GST-agarose beads after thrombin cleavage; 8: WalK-PAS^FULL^ eluted after thrombin cleavage of fusion protein; 9: monomer peak resulting from SEC purification of WalK-PAS^FULL^. **(B)** Chromatogram from SEC purification of WalK-PAS^FULL^ showing the presence of both monomeric proteins and large aggregates; **(C)** Chromatogram obtained by re-chromatographing the monomer peak from panel B.

**Suppl Fig 2. Incorporation of selenomethionine in WalK-PAS^FULL^and detection by MALDI-MS. (A)**

Amino acids from 251-376 of the WalK-PAS^FULL^ construct with methionine residues highlighted in red. (B) Peptide fragmentation prediction for WalK-PAS^FULL^ for trypsin digestion and the detected 24 fragments. The highlighted corrected masses were detected using MALDI-MS. **(C)** MALDI spectra of tryptic fragments showing the corrected mass (highlighted in purple). The molecular weight difference in the individual fragmented peptide corresponded to the number of selenomethionines in the peptide. The mass analysis verified the 100% substitution of SeMet in WalK-PAS^FUL^ domain.

**Suppl Fig 3. Structural comparison of WalK-PAS^FULL^ with WalK-PAS^TRUNC^.** (A) The Zn^2+^ - coordinating residues of WalK-PAS^FULL^ are shown as cyan sticks, with the atoms contributing to the interactions as spheres. The coordinating bonds are illustrated with black dashed lines. The electron density shown is the 2Fº-F_C_ map contoured at 1.5 σ. **(B)** Superposition of the crystal structure of WalK-PAS^FULL^ (green) with WalK-PAS^TRUNC^ (light blue) in cartoon representation. The bound Zn^2+^ ions are shown as spheres and the Zn^2+^-coordinating residues are shown as sticks. **(B)** The Zn^2+^-binding site of WalK-PAS^TRUNC^as described in (A).

**Suppl Fig 4. Homology modelling map of pdb structures used to build the 5. *aureus* WalK model.**

Red region corresponds to structure pdb 5IS1 (residues 33-182, identity 100%), orange to pdb structure 4MN6 (residues 266 to 371, identity 100%), and underlined to pdb structure 4I5S (residues 223 to 599, identity 44%). Blue regions did not have significant similarities to the pdb. The N-terminus (1-32) was modelled as transmembrane helix, as was the middle sequence (183-222). The C-terminus was modelled as disordered.

## References

1. Tong, S. Y., Davis, J. S., Eichenberger, E., Holland, T. L. & Fowler, V. G., Jr. Staphylococcus aureus infections: epidemiology, pathophysiology, clinical manifestations, and management. Clin Microbiol Rev 28, 603–661, doi:10.1128/CMR.00134-14 (2015).

2. Howden, B. P., Davies, J. K., Johnson, P. D., Stinear, T. P. & Grayson, M. L. Reduced vancomycin susceptibility in Staphylococcus aureus, including vancomycin-intermediate and heterogeneous vancomycin-intermediate strains: resistance mechanisms, laboratory detection, and clinical implications. Clin Microbiol Rev 23, 99–139, doi:10.1128/CMR.00042- 09 (2010).

3. Kelley, P. G., Gao, W., Ward, P. B. & Howden, B. P. Daptomycin non-susceptibility in vancomycin-intermediate Staphylococcus aureus (VISA) and heterogeneous-VISA (hVISA): implications for therapy after vancomycin treatment failure. J Antimicrob Chemother 66, 1057–1060, doi:10.1093/jac/dkr066 (2011).

4. Chua, K. & Howden, B. P. Treating Gram-positive infections: vancomycin update and the whys, wherefores and evidence base for continuous infusion of anti-Gram-positive antibiotics. Curr Infect Dis Rep 22, 525–534, doi:10.1097/QCO.0b013e328331fbcd (2009).

5. Holmes, N. E. et al. Antibiotic Choice May Not Explain Poorer Outcomes in Patients With Staphylococcus aureus Bacteremia and High Vancomycin Minimum Inhibitory Concentrations. J Infect Dis 204, 340–347, doi:10.1093/infdis/jir270 (2011).

6. Dubrac, S., Bisicchia, P., Devine, K. M. & Msadek, T. A matter of life and death: cell wall homeostasis and the WalKR (YycGF) essential signal transduction pathway. Mol Microbiol 70, 1307–1322, doi:10.1111/j.1365-2958.2008.06483.x (2008).

7. Bem, A. E. et al. Bacterial Histidine Kinases as Novel Antibacterial Drug Targets. Acs Chem Biol 10, 213–224, doi:10.1021/cb5007135 (2015).

8. Fabret, C. & Hoch, J. A. A two-component signal transduction system essential for growth of Bacillus subtilis: implications for anti-infective therapy. J Bacteriol 180, 6375–6383 (1998).

9. Martin, P. K., Li, T., Sun, D., Biek, D. P. & Schmid, M. B. Role in cell permeability of an essential two-component system in Staphylococcus aureus. J Bacteriol 181, 3666–3673 (1999).

10. Dubrac, S. & Msadek, T. Identification of genes controlled by the essential YycG/YycF two-component system of Staphylococcus aureus. J Bacteriol 186, 1175–1181 (2004).

11. Howell, A. et al. Genes controlled by the essential YycG/YycF two-component system of Bacillus subtilis revealed through a novel hybrid regulator approach. Mol Microbiol 49, 1639–1655, doi:10.1046/j.1365-2958.2003.03661.x (2003).

12. Hancock, L. E. & Perego, M. Systematic inactivation and phenotypic characterization of two-component signal transduction systems of Enterococcus faecalis V583. J Bacteriol 186, 7951–7958, doi:10.1128/JB.186.23.7951-7958.2004 (2004).

13. Kallipolitis, B. H. & Ingmer, H. Listeria monocytogenes response regulators important for stress tolerance and pathogenesis. FEMS Microbiol Lett 204, 111–115 (2001).

14. Villanueva, M. et al. Sensory deprivation in Staphylococcus aureus. Nat Commun 9, doi:10.1038/s41467-018-02949-y (2018).

15. Fukuchi, K. et al. The essential two-component regulatory system encoded by yycF and yycG modulates expression of the ftsAZ operon in Bacillus subtilis. Microbiology 146 (Pt 7), 1573–1583, doi:10.1099/00221287-146-7-1573 (2000).

16. Dubrac, S., Boneca, I. G., Poupel, O. & Msadek, T. New insights into the WalK/WalR (YycG/YycF) essential signal transduction pathway reveal a major role in controlling cell wall metabolism and biofilm formation in Staphylococcus aureus. J Bacteriol 189, 8257–8269, doi:10.1128/JB.00645-07 (2007).

17. Howden, B. P. et al. Genomic analysis reveals a point mutation in the two-component sensor gene graS that leads to intermediate vancomycin resistance in clinical Staphylococcus aureus. Antimicrob Agents Ch 52, 3755–3762, doi:10.1128/Aac.01613-07 (2008).

18. Mwangi, M. M. et al. Tracking the in vivo evolution of multidrug resistance in Staphylococcus aureus by whole-genome sequencing. Proc Natl Acad Sci U S A 104, 9451–9456, doi:10.1073/pnas.0609839104 (2007).

19. Neoh, H. M. et al. Mutated response regulator graR is responsible for phenotypic conversion of Staphylococcus aureus from heterogeneous vancomycin-intermediate resistance to vancomycin-intermediate resistance. Antimicrob Agents Ch 52, 45–53, doi:10.1128/Aac.00534-07 (2008).

20. Hu, Q., Peng, H. & Rao, X. Molecular Events for Promotion of Vancomycin Resistance in Vancomycin Intermediate Staphylococcus aureus. Front Microbiol 7, 1601, doi:10.3389/fmicb.2016.01601 (2016).

21. Shoji, M. et al. walK and clpP Mutations Confer Reduced Vancomycin Susceptibility in Staphylococcus aureus. Antimicrob Agents Ch 55, 3870–3881, doi:10.1128/Aac.01563-10 (2011).

22. Howden, B. P. et al. Evolution of Multidrug Resistance during Staphylococcus aureus Infection Involves Mutation of the Essential Two Component Regulator WalKR. Plos Pathogens 7, doi:10.1371/journal.ppat.1002359 (2011).

23. Hu, J. F., Zhang, X., Liu, X. Y., Chen, C. & Sun, B. L. Mechanism of Reduced Vancomycin Susceptibility Conferred by walK Mutation in Community-Acquired Methicillin-Resistant Staphylococcus aureus Strain MW2. Antimicrob Agents Ch 59, 1352–1355, doi:10.1128/Aac.04290-14 (2015).

24. Fukushima, T. et al. A role for the essential YycG sensor histidine kinase in sensing cell division. Mol Microbiol 79, 503–522, doi:10.1111/j.1365-2958.2010.07464.x (2011).

25. Bisicchia, P. et al. The essential YycFG two-component system controls cell wall metabolism in Bacillus subtilis. Mol Microbiol 65, 180–200, doi:10.1111/j.1365-2958.2007.05782.x (2007).

26. Delaune, A. et al. The WalKR System Controls Major Staphylococcal Virulence Genes and Is Involved in Triggering the Host Inflammatory Response. Infect Immun 80, 3438–3453, doi:10.1128/Iai.00195-12 (2012).

27. Delaune, A. et al. Peptidoglycan crosslinking relaxation plays an important role in Staphylococcus aureus WalKR-dependent cell viability. Plos One 6, e17054, doi:10.1371/journal.pone.0017054 (2011).

28. Cameron, D. R., Jiang, J. H., Kostoulias, X., Foxwell, D. J. & Peleg, A. Y. Vancomycin susceptibility in methicillin-resistant Staphylococcus aureus is mediated by YycHI activation of the WalRK essential two-component regulatory system. Sci Rep-Uk 6, doi:10.1038/srep30823 (2016).

29. Szurmant, H., Mohan, M. A., Imus, P. M. & Hoch, J. A. YycH and YycI interact to regulate the essential YycFG two-component system in Bacillus subtilis. J Bacteriol 189, 3280–3289, doi:10.1128/JB.01936-06 (2007).

30. Henry, J. T. & Crosson, S. Ligand-Binding PAS Domains in a Genomic, Cellular, and Structural Context. Annu Rev Microbiol 65, 261–286, doi:10.1146/annurev-micro-121809-151631 (2011).

31. Poupel, O. et al. Transcriptional Analysis and Subcellular Protein Localization Reveal Specific Features of the Essential WalKR System in Staphylococcus aureus. Plos One 11, e0151449, doi:10.1371/journal.pone.0151449 (2016).

32. Monk, I. R., Howden, B. P., Seemann, T. & Stinear, T. P. Correspondence: Spontaneous secondary mutations confound analysis of the essential two-component system WalKR in Staphylococcus aureus. Nat Commun 8, doi:10.1038/ncomms14403 (2017).

33. Pasztor, L. et al. Staphylococcal Major Autolysin (Atl) Is Involved in Excretion of Cytoplasmic Proteins. J Biol Chem 285, 36794–36803, doi:10.1074/jbc.M110.167312 (2010).

34. Bose, J. L., Lehman, M. K., Fey, P. D. & Bayles, K. W. Contribution of the Staphylococcus aureus Atl AM and GL Murein Hydrolase Activities in Cell Division, Autolysis, and Biofilm Formation. Plos One 7, doi:10.1371/journal.pone.0042244 (2012).

35. Gong, W. M. et al. Structure of a biological oxygen sensor: A new mechanism for heme-driven signal transduction. P Natl Acad Sci USA 95, 15177–15182, doi:DOI 10.1073/pnas.95.26.15177 (1998).

36. Laitaoja, M., Valjakka, J. & Janis, J. Zinc coordination spheres in protein structures. Inorg Chem 52, 10983–10991, doi:10.1021/ic401072d (2013).

37. Hardt, P. et al. The cell wall precursor lipid II acts as a molecular signal for the Ser/Thr kinase PknB of Staphylococcus aureus. Int J Med Microbiol 307, 1–10, doi:10.1016/j.ijmm.2016.12.001 (2017).

38. Libby, E. A., Goss, L. A. & Dworkin, J. The Eukaryotic-Like Ser/Thr Kinase PrkC Regulates the Essential WalRK Two-Component System in Bacillus subtilis. Plos Genet 11, doi:10.1371/journal.pgen.1005275 (2015).

39. Wang, C. et al. Mechanistic Insights Revealed by the Crystal Structure of a Histidine Kinase with Signal Transducer and Sensor Domains. Plos Biol 11, doi:10.1371/journal.pbio.1001493 (2013).

40. Jacob-Dubuisson, F., Mechaly, A., Betton, J. M. & Antoine, R. Structural insights into the signalling mechanisms of two-component systems. Nat Rev Microbiol, doi:10.1038/s41579-018-0055-7 (2018).

41. Mullner, M. et al. A PAS Domain with an Oxygen Labile [4Fe-4S](2+) Cluster in the Oxygen Sensor Kinase NreB of Staphylococcus carnosus. Biochemistry-Us 47, 13921–13932, doi:10.1021/bi8014086 (2008).

42. Little, R., Martinez-Argudo, I. & Dixon, R. Role of the central region of NifL in conformational switches that regulate nitrogen fixation. Biochem Soc T 34, 162–164, doi:10.1042/Bst0340162 (2006).

43. Cheung, J., Bingman, C. A., Reyngold, M., Hendrickson, W. A. & Waldburger, C. D. Crystal structure of a functional dimer of the PhoQ sensor domain. J Biol Chem 283, 13762–13770, doi:10.1074/jbc.M710592200 (2008).

44. Grundling, A., Missiakas, D. M. & Schneewind, O. Staphylococcus aureus mutants with increased lysostaphin resistance. J Bacteriol 188, 6286–6297, doi:10.1128/Jb.00457-06 (2006).

45. Poupel, O., Proux, C., Jagla, B., Msadek, T. & Dubrac, S. SpdC, a novel virulence factor, controls histidine kinase activity in Staphylococcus aureus. PLoS Pathog 14, e1006917, doi:10.1371/journal.ppat.1006917 (2018).

46. Echenique, J. R. & Trombe, M. C. Competence repression under oxygen limitation through the two-component MicAB signal-transducing system in Streptococcus pneumoniae and involvement of the PAS domain of MicB. J Bacteriol 183, 4599–4608, doi:Doi 10.1128/Jb.183.15.4599-4608.2001 (2001).

47. Zhang, Y., Werling, U. & Edelmann, W. SLiCE: a novel bacterial cell extract-based DNA cloning method. Nucleic Acids Res 40, e55, doi:10.1093/nar/gkr1288 (2012).

48. Monk, I. R., Tree, J. J., Howden, B. P., Stinear, T. P. & Foster, T. J. Complete Bypass of Restriction Systems for Major Staphylococcus aureus Lineages. Mbio 6, doi:10.1128/mBio.00308-15 (2015).

49. Monk, I. R., Shah, I. M., Xu, M., Tan, M. W. & Foster, T. J. Transforming the Untransformable: Application of Direct Transformation To Manipulate Genetically Staphylococcus aureus and Staphylococcus epidermidis. Mbio 3, doi:10.1128/mBio.00277-11 (2012).

50. Lee, J. Y. H. et al. Global spread of three multidrug-resistant lineages of Staphylococcus epidermidis. Nat Microbiol, doi:10.1038/s41564-018-0230-7 (2018).

51. Monk, I. R., Cook, G. M., Monk, B. C. & Bremer, P. J. Morphotypic conversion in Listeria monocytogenes biofilm formation: biological significance of rough colony isolates. Appl Environ Microbiol 70, 6686–6694, doi:10.1128/AEM.70.11.6686-6694.2004 (2004).

52. McCoy, A. J. et al. Phaser crystallographic software. J Appl Crystallogr 40, 658–674, doi:10.1107/S0021889807021206 (2007).

53. Evans, P. R. An introduction to data reduction: space-group determination, scaling and intensity statistics. Acta Crystallogr D Biol Crystallogr 67, 282–292, doi:10.1107/S090744491003982X (2011).

54. Adams, P. D. et al. PHENIX: a comprehensive Python-based system for macromolecular structure solution. Acta Crystallogr D Biol Crystallogr 66, 213–221, doi:10.1107/S0907444909052925 (2010).

55. Emsley, P., Lohkamp, B., Scott, W. G. & Cowtan, K. Features and development of Coot. Acta Crystallogr D Biol Crystallogr 66, 486–501, doi:10.1107/S0907444910007493 (2010).

56. Humphrey, W., Dalke, A. & Schulten, K. VMD: visual molecular dynamics. J Mol Graph 14, 33-38, 27-38 (1996).

57. Kim, T. et al. Structural Studies on the Extracellular Domain of Sensor Histidine Kinase YycG from Staphylococcus aureus and Its Functional Implications. J Mol Biol 428, 3074–3089, doi:10.1016/j.jmb.2016.06.019 (2016).

58. Phillips, J. C. et al. Scalable molecular dynamics with NAMD. J Comput Chem 26, 1781–1802, doi:10.1002/jcc.20289 (2005).

59. Brooks, B. R. et al. CHARMM: the biomolecular simulation program. J Comput Chem 30, 1545–1614, doi:10.1002/jcc.21287 (2009).

60. Stivala, A., Wybrow, M., Wirth, A., Whisstock, J. C. & Stuckey, P. J. Automatic generation of protein structure cartoons with Pro-origami. Bioinformatics 27, 3315–3316, doi:10.1093/bioinformatics/btr575 (2011).

61. Baker, N. A., Sept, D., Joseph, S., Holst, M. J. & McCammon, J. A. Electrostatics of nanosystems: application to microtubules and the ribosome. Proc Natl Acad Sci U S A 98, 10037–10041, doi:10.1073/pnas.181342398 (2001).

